# tWhEEL: a transcript-aware workflow for quantifying CRISPR knockout feasibility using transcriptome assembly

**DOI:** 10.64898/2026.10.27.762389

**Authors:** Kazuki Nakamae, Hidemasa Bono

**Author notes:** Corresponding authors (H.B); (KN).

## Abstract

Gene redundancy can complicate loss-of-function experiments (knockout), and its extent varies among organisms and biological functions. Relating variation in gene redundancy to the availability of genome-editing targets can help researchers select tractable biological functions and experimental organisms. Here, we present the transcript-aware Whole-genomic Evaluation for Editing Locus (tWhEEL), a pipeline that infers redundancy in silico from transcript information and cross-references it with specific guide RNA designs. tWhEEL integrates CRISPR-Cas9 guide design with Gene Ontology (GO) and pathway annotations to compute a knockout feasibility index (KFI) for individual genes and functional categories. Interactive HTML reports link GO-associated KFI and pathway-associated KFI summaries to the contributing groups, genes, and guide candidates with links to QuickGO or WikiPathways. By linking sequence-level guide candidates to functional-level targetability, tWhEEL can support the selection of biological functions and organisms for reverse-genetic experiments. Applications to model and nonmodel datasets revealed distinct targetability profiles, including high full targetability in yeast and query-protein-set dependence in zebrafish. A zebrafish Fanconi anemia case study recovered 14 of 19 published target-site designs. By connecting sequence-level guide candidates with function-level experimental planning, tWhEEL provides a framework for comparing reverse genetic tractability and prioritizing biological functions and organisms for further study.

## INTRODUCTION

The clustered regularly interspaced short palindromic repeats (CRISPR)-associated protein 9 (Cas9) system has made targeted reverse genetics feasible in a broad range of organisms(1–13); however, the practical interpretation of gene knockout depends on factors beyond the presence of a protospacer adjacent motif (PAM) and a seemingly specific guide RNA. Genes frequently belong to duplicated families, retain partially redundant paralogs, or are represented by multiple transcript isoforms. In polyploid or recently duplicated genomes, the same experimental perturbation may need to remove several homologous loci to produce a measurable phenotype, as previously reported in studies targeting homologous genes in wheat (14). Gene complexity has frequently been discussed in nonmodel organisms. For example, at the gene-group level, adaptive diversification of the HMA4 region has resulted in three tandem copies in *Arabidopsis halleri* (15), whereas in *Arabidopsis thaliana*, this region has one copy. Conversely, the absence of redundant copies can render a nonmodel organism particularly informative for functional genetics. Examples include the single-copy phototropin gene system in *Marchantia polymorpha*, where a reduced redundancy architecture has helped clarify photoreceptor function (16). These considerations are frequently treated qualitatively in the design of knockout experiments. Gene families may be inspected to assess whether the proposed knockout is likely to be interpretable, followed by the design of one or several single-guide RNAs. However, no standard framework has been widely adopted for quantifying whether gene groups in an active transcriptome can be operationally targeted through genome editing at the level of functional annotations or pathways.

Existing tools such as CRISPOR and CHOPCHOP support guide selection through sequence-specificity assessment, guide ranking, and, where annotations are available, consideration of transcript isoforms (17, 18). Experimental planning relates guide RNA candidates to a set of annotated genes associated with specific biological functions using the software. The design strategy is, however, limited to humans and major model organisms because gene models and functional resources remain incomplete in most organisms. We suppose that although the database for the focus species is incomplete, an explicit summary of guide availability in functional genes can help identify which functional gene targets are represented and whether functional knockout design is feasible for the phenotype focused on by the researcher.

Here, we describe the transcript-aware Whole-genomic Evaluation for Editing Locus (tWhEEL) pipeline, a transcript-aware workflow for genome-editing target search and knockout feasibility evaluation across functional gene groups. tWhEEL builds on the functional annotation approach of the FanFlow transcriptome annotation workflows (19), gene-group classification, CRISPR guide design, and functional summarization. The key output is the Knockout Feasibility Index (KFI), a simple annotation-level score that quantifies the fraction of units associated with a biological function, Gene Ontology (GO) term (20), or pathway for which at least one suitable sgRNA can be designed. By coupling guide design with functional annotations, tWhEEL aims to provide both target-level guide candidates and organism-level summaries to inform the experimental field across various organisms and support genome editing design leading to knockout phenotype analysis.

## MATERIALS AND METHODS

### Workflow implementation and input modes

tWhEEL is a Python workflow that combines protein annotation, transcript-derived guide discovery, sequence-specificity filtering, and functional summaries (Figure 1A). For the benchmarks reported here, guide design and GO-KFI and PW-KFI calculations were performed using tWhEEL v1.3.4. The corresponding HTML reports were subsequently regenerated using tWhEEL v1.3.5, which corrected the display of GO-KFI category counts and stacked bars. This report regeneration did not repeat guide design or KFI calculation and left the underlying analysis results unchanged. The analysis settings and input paths are supplied in a YAML configuration file. The workflow supports two input modes: protein prediction from a transcriptome assembly or analysis of supplied protein sequences, together with transcriptome and protein–transcript mapping. In the latter mode, mapping is a headerless table with transcript identifiers in the first column and protein identifiers in the second column. Each protein must be mapped unambiguously to a non-empty transcript record, and mapping protein identifiers must match those in the supplied protein set. These requirements were validated before functional annotation. Genome assembly can be used as a reference for sequence specificity assessments. The detailed process is summarized as follows.

**Figure 1:**
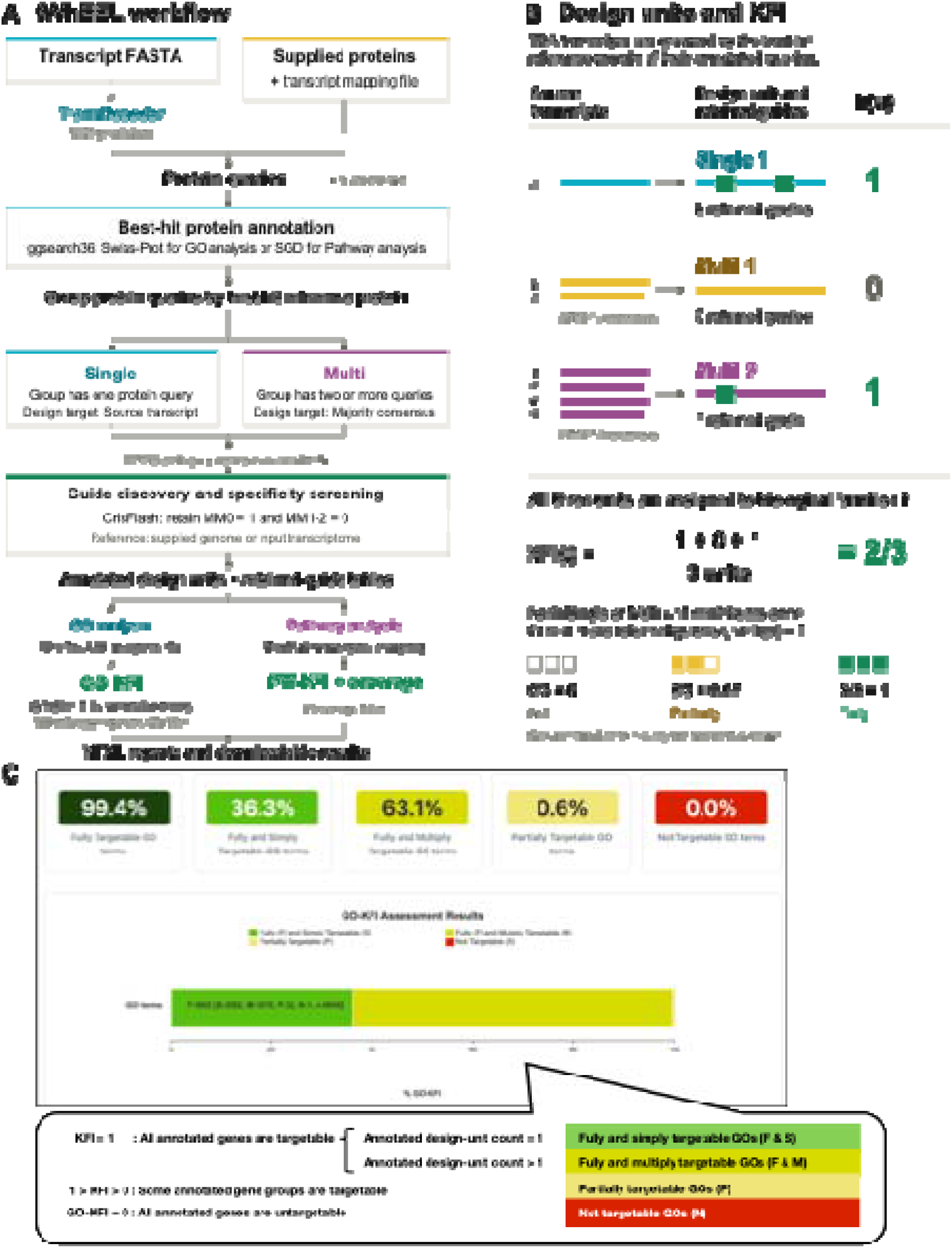
The pipeline overview of tWhEEL. (A) Overview of the tWhEEL workflow. Protein queries are predicted from input transcripts using TransDecoder or supplied with a protein-to-transcript mapping; transcript FASTA is required in both routes. BUSCO assesses protein-set completeness but does not filter queries for downstream analysis. Depending on the selected profile, ggsearch36 assigns best-hit reference proteins from Swiss-Prot for GO analysis or SGD for the yeast pathway analysis. Queries sharing a reference protein are grouped and mapped back to their whole source transcripts. Single groups contain one query, whereas multigroups contain two or more queries; these counts do not necessarily correspond to distinct genes or transcripts. Each single group supplies its source transcript, and each multigroup supplies a representative sequence generated by MAFFT alignment and nongap majority consensus. IUPAC ambiguity codes are converted to N before guide discovery. CrisFlash screens transcript-derived candidates against a supplied genome or, if absent, the input transcriptome, retaining candidates with exactly one perfect match and no matches with one or two mismatches. The GO profile retrieves UniProt GO assignments and calculates GO-KFI; terms with GO-KFI = 1 are subsequently summarized using the GO ontology and generic GO Slim. The alternative yeast pathway profile maps annotated units to WikiPathways genes, calculates PW-KFI and annotation coverage, and applies the configured coverage filter. Both profiles produce HTML reports and downloadable results. (B) Design units and calculation of the Knockout Feasibility Index (KFI). Seven illustrative TSA transcripts, each represented by one annotated protein query, are grouped by their best-hit reference protein into one single and two multi design units. Single units retain their source transcript sequence, whereas multi-units use an alignment-derived representative consensus. Green blocks denote retained guide sites. Each unit contributes h(u) = 1 if it has at least one retained guide and h(u) = 0 otherwise. With all three units assigned to biological function t, guide counts of 2, 0, and 1 give KFI(t) = (1 + 0 + 1)/3 = 2/3. Each unit contributes once, independently of its transcript or guide count. Alternative guide-availability states illustrate fully, partially, and not targetable functions. Sequence bars are not to scale. (C) The screenshot of the HTML report describing the gene ontology-based KFI value. The yeast GO-KFI distribution and four targetability classes are displayed. The terms with KFI = 1, which indicates all annotated genes are targetable, are subsequently summarized into “Fully Targetable (F)”. The F label is split into two subclasses: “Fully and Simply Targetable (F&S)” and “Fully and Multiply Targetable (F&M).” F&S denotes GO terms containing one annotated design unit with at least one retained guide, whereas F&M denotes GO terms containing multiple annotated design units, each with at least one retained guide. Terms with KFI between 0 and 1 are classified as “Partially targetable (P),” indicating that the guide design can be eligible for a part of the annotated gene. When all annotated genes are untargetable, the KFI is zero, and the terms with KFI = 0 are classified into “Not targetable GOs (N).” The classes of F, F&S, F&M, P, and N are illustrated as color boxes with dark green, light green, dark yellow, yellow, and red, respectively. The percentage is rounded to one decimal place. The exact counts are shown in the stacked bar chart below. Each count of the classified terms is shown with the total value “n.” The statistics are shortly summarized as “F:5622 [S:2052, M:3570, P:32, N:1, n:5655].” The GO-KFI analysis used tWhEEL v1.3.4, and the HTML report shown here was regenerated using v1.3.5 without rerunning guide design or GO-KFI calculation.

### 1. Protein prediction and sequence assessment

For Transcriptome Shotgun Assembly (TSA) inputs, the coding regions were predicted using TransDecoder.LongOrfs (v5.7.1) (21) with a minimum protein-length setting of 100 amino acids, followed by TransDecoder.Predict (v5.7.1) (21). The length setting applied to the LongOrfs step was not used as a separate filter for every final predicted protein. When the user provides protein–transcript mapping, tWhEEL skips the above protein prediction step. Protein sequence statistics were obtained using SeqKit (v2.8.0) (22), and the protein set completeness of TSA was assessed using BUSCO (v5.2.2) (23) in the protein mode.

### 2. Homology-based functional annotation and gene-group classification

Protein sequences were compared with the reference protein database using ggsearch36 (v. 36.3.8g) global alignment from the FASTA suite (24). Searches used an E-value threshold of 0.1 and retained the top-ranked hit for each annotated query, as previously reported in the FanFlow pipeline. FanFlow annotation parses the best-hit subject identifiers, descriptions, and gene name fields when available. The UniProtKB/Swiss-Prot database (25) must be used as the reference protein database when downstream analysis includes GO analysis. The Saccharomyces Genome Database (SGD) (26) was used for downstream pathway analysis.

The annotated queries were grouped according to their best-hit reference protein identifiers. A group containing one query was assigned a “Single” identifier; a group containing multiple queries was assigned a “multi” identifier. Each identifier defined an annotation-based design unit. These units do not establish the genomic copy number, orthology, or functional redundancy because multiple queries can reflect transcript isoforms, homologous loci, or assembly redundancy.

### 3. Multigroup alignment and consensus generation

The annotated proteins were linked to their source transcripts using the supplied mapping or transcript identifier encoded in the TransDecoder protein identifier. Guide discovery used complete transcript records, rather than sequences restricted to predicted coding regions. Single units retained their corresponding transcript sequences. For multi-units, the member transcripts were aligned using MAFFT (v7.505) (27). Each MAFFT process used one thread, and the number of concurrent group alignments was controlled separately by the workflow configuration.

For groups containing more than one sequence, a consensus sequence was generated by taking the most frequent nongap nucleotides at each alignment position (Supplementary Figure 1A). Ties were resolved by the first character encountered in the aligned records, and columns containing only gaps were represented by N. The consensus sequence was then used as a representative target sequence for guide design, enabling tWhEEL to search for guide candidates that can address grouped transcripts or homolog sets. The queries with “single” identifier, single-copy-like groups, were processed without the above consensus generation step. Finally, some IUPAC ambiguity codes (28) other than N, which were not supported by downstream software, were converted to N in the specificity reference sequences, single sequences, and integrated multi-consensus sequences before the downstream guide design step.

### 4. CRISPR-Cas9 guide design and sequence-specificity filtering

The CRISPR guide design step with Crisflash (v1.2.0) (29) searched the prepared transcript-derived sequences for candidate sites and assessed matches in the selected specificity reference with a maximum of two mismatches with NGG-3′ PAM (Supplementary Figure 1B). A candidate was retained only when it had exactly one perfect match and no matches with one or two mismatches in the reference, corresponding to MM0 = 1 and MM1–2 = 0. The benchmark configurations used 10 threads, and the memory limits were specified separately for each analysis.

If a genome assembly is available, it is used as a reference for off-target searches. If no genome assembly is provided, tWhEEL returns to transcriptome-derived sequences, which is useful for early stage nonmodel organisms but less comprehensive for genome-wide off-target evaluation. Even in genome-backed runs, the filter establishes the number of reference sequence matches without independently verifying whether the retained genomic match belongs to the intended locus or coding exon.

The Crisflash outputs were filtered to retain the target-specific guide candidates according to the above criteria. The guide information was merged back into the functional annotation table as a guide_ target field, preserving the group identifiers, guide sequences, coordinates, strands, and off-target summary counts.

### 5. Knockout Feasibility Index

tWhEEL assessed the feasibility of gene knockout using the KFI for each biological function in the target organisms and tissues (Figure 1B). The index was calculated using the above annotation table and guide design collection. The KFI connects genes or gene groups associated with biological functions to the availability of suitable sgRNAs (Figure 1B). For a biological function t, let U(t) denote its counting units: distinct annotation-based design units assigned to a GO term, or evaluable canonical genes represented in a pathway. Let h(u) be equal to 1 when unit u has at least one retained guide, and 0 otherwise. KFI is defined as:

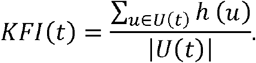

KFI ranges from 0 to 1. Values of 1, between 0 and 1, and 0 indicate fully, partially, and not targetable functions, respectively. Fully targetable functions are further classified as fully and simply targetable when represented by one counting unit, and as fully and multiply targetable when represented by multiple counting units. The classification distinguishes functions represented by one targetable unit from those for which several units are targetable, linking functional grouping to experimental planning and providing practical insights into gene functional redundancy for genome editing.

### 5-A. GO analysis and GO-KFI report generation

GO annotations were retrieved from UniProt accessions through the UniProt REST API using the biological process, molecular function, and cellular component fields and were summarized in tWhEEL output tables. tWhEEL produced GO-annotated guide design tables, computed the KFI of GO terms (named GO-KFI) (Supplementary Figure 1C), generated GO-level summary statistics, and created a reverse-lookup table linking each GO term to the contributing tWhEEL gene groups and guide RNAs. GO Slim parent-term summaries were generated using GO resources from the GO knowledgebase and GOATOOLS. The HTML report includes summary cards, searchable GO-KFI tables, QuickGO links, and downloadable guide RNA tables for individual GO terms.

### 5-B. *Pathway analysis and pathway-associated KFI report generation*

Pathway analysis uses WikiPathways Graph Pathway Markup Language (GPML) files, which are open and community-curated pathway resources (30). In the current tWhEEL version (v1.3.5), pathway-gene matching is implemented for SGD identifiers and Ensembl systematic identifiers and is therefore the most mature for yeast datasets. Pathway identifiers were matched exactly to the annotation records using cross-references from the configured SGD FASTA headers to connect systematic identifiers to SGD gene identities. The repeated nodes and cross-database aliases resolving to the same canonical genes were counted as the same. tWhEEL calculated the KFI value for each pathway, named pathway-associated (PW)-KFI (Supplementary Figure 1D), and produced pathway- and gene-level output tables (Figure 3B). Pathways without evaluable genes or unresolved cross-database canonical identities were not assigned a score. The annotation coverage was calculated as the proportion of evaluable canonical pathway genes that matched the annotation records. Pathway-associated KFI (PW-KFI) used the same denominator and counted genes with at least one guide retained in the numerator. The analysis was configured to exclude pathways with annotation coverage below 0.5. The interactive pathway report visualizes GPML-derived pathway diagrams and highlights genes according to guide availability. Clicking on a pathway gene provides guide-level details, including guide sequences and off-target summary fields.

### Datasets and analysis settings for benchmarking

The transcriptome comparison comprised eight TSA datasets from the NCBI GenBank database: *Homo sapiens* (GenBank: GLNK00000000.1), *Danio rerio* (GenBank: GFIL00000000.1) (31), *Drosophila melanogaster* (GenBank: GHHI00000000.1), *Arabidopsis thaliana* (GenBank: GKIF00000000.1) (32), *Saccharomyces cerevisiae* Y12 (GenBank: GFJR00000000.1), *Nannochloropsis gaditana* (GenBank: GAGR00000000.1), *Mimachlamys varia* (GenBank: GGGO00000000.1) (33) and *Entoria okinawaensis* (GenBank: ICSG00000000.1) (34). The analyses used predicted proteins and input transcriptomes as specificity references; no genome assemblies were used for this comparison.

*Saccharomyces cerevisiae* CEN.PK113-7D analyses used cDNA and protein sequences from the Ensembl Fungi release 62, with the corresponding repeat-masked ASM26988v1 genome assembly (35). Separate GO and pathway analyses were performed for protein annotation using UniProtKB/Swiss-Prot and SGD S288C reference protein sets, respectively. The analysis of the CEN.PK113-7D strain was distinct from that of the Y12 strain (Table 1).

**Table 1:** GO-KFI profiles across eight model and nonmodel transcriptome datasets.

| Organism | TSA (GenBank accession NO.) | Complete gene rate reported by Protein BUSCO (Database) | Fully and simply targetable GOs [%] | Fully and multiply targetable [%] | Partially targetable[%] | Not targetable[%] | Total GOs |
|---|---|---|---|---|---|---|---|
| *Homo sapiens* | GLNK00000000.1 | 82.0% (primates_odb10) | 29.1 | 66.4 | 4.5 | 0.1 | 17858 |
| *Danio rerio* | GFIL00000000.1 | 94.7% (actinopterygii_odb10) | 27.1 | 65.1 | 7.6 | 0.2 | 17664 |
| *Drosophila melanogaster* | GHHI00000000.1 | 90.3% (diptera_odb10) | 29.4 | 24.7 | 37.6 | 8.4 | 13227 |
| *Arabidopsis thaliana* | GKIF00000000.1 | 98.9% (brassicales_odb10) | 18.8 | 11.1 | 54.2 | 15.8 | 14146 |
| *Saccharomyces cerevisiae* Y12 | GFJR00000000.1 | 89.0% (saccharomycetes_odb10) | 37.6 | 52.6 | 6.4 | 3.4 | 5884 |
| *Nannochloropsis gaditana* | GAGR00000000.1 | 75.0% (stramenopiles_odb10) | 46.6 | 53.4 | 0 | 0 | 10174 |
| *Mimachlamys varia* | GGGO00000000.1 | 90.2% (mollusca_odb10) | 19.9 | 14.6 | 52.8 | 12.7 | 17116 |
| *Entoria okinawaensis* | ICSG00000000.1 | 96.5% (insecta_odb10) | 23.6 | 17.1 | 48.1 | 11.1 | 15377 |

Zebrafish RefSeq analyses used the GRCz12tu assembly (GCF_049306965.1) (36), its transcript sequences, and two matched query protein sets: an NP-only set and a set containing NP-and XP-prefixed records. These prefixes distinguish known from model RefSeq proteins. Protein entries without resolvable transcript counterparts were excluded during input preparation. The NP-only and NP+XP inputs contained 16,542 and 89,814 protein–transcript pairs, respectively. Both analyses used the same UniProtKB/Swiss-Prot annotation database and the same genomic specificity reference. Thus, the comparison changed the query protein set rather than the reference database used for homology-based functional annotation.

For GO analyses, the local UniProtKB/Swiss-Prot FASTA was used as the homology-search database. Yeast pathway analysis was performed using the WikiPathways *Saccharomyces cerevisiae* GPML collection, dated September 10, 2025. Exact sequence accessions, protein set filters, configuration files, and database snapshots should accompany the deposited analysis data. The release or retrieval dates and checksums of the Swiss-Prot FASTA, SGD FASTA, GO ontology, and generic GO Slim files are listed in Supplementary Table 1.

The BUSCO lineages were primates_odb10, actinopterygii_odb10, diptera_odb10, brassicales_odb10, saccharomycetes_odb10, stramenopiles_odb10, mollusca_odb10, and insecta_odb10 (Table 1). The CEN.PK113-7D and zebrafish RefSeq analyses used saccharomycetes_odb10 and actinopterygii_odb10, respectively. The reference resource versions and BUSCO lineage dataset are listed in Supplementary Table 1.

### Published zebrafish target-site comparison

Zebrafish RefSeq analyses were compared using NP-only and NP+XP query-protein sets to assess the effect of including predicted protein records. The NP-only GRCz12tu analysis was compared with the published target sites for 19 zebrafish Fanconi anemia (FA) pathway genes reported by Ramanagoudr-Bhojappa et al (37). Table 2 shows the representation in the protein annotation, presence of the published site in the genome, eligible mapped transcripts, passage through the sequence-specificity filter, and retention in the final output. Workflow stages lacking eligible upstream proteins or sequences were recorded as not evaluated. GO:0043240 annotations were recorded separately from the guide recovery. The first nucleotide of the NGG PAM sequence is represented as N for a consistent display of published target-site sequences.

**Table 2:**
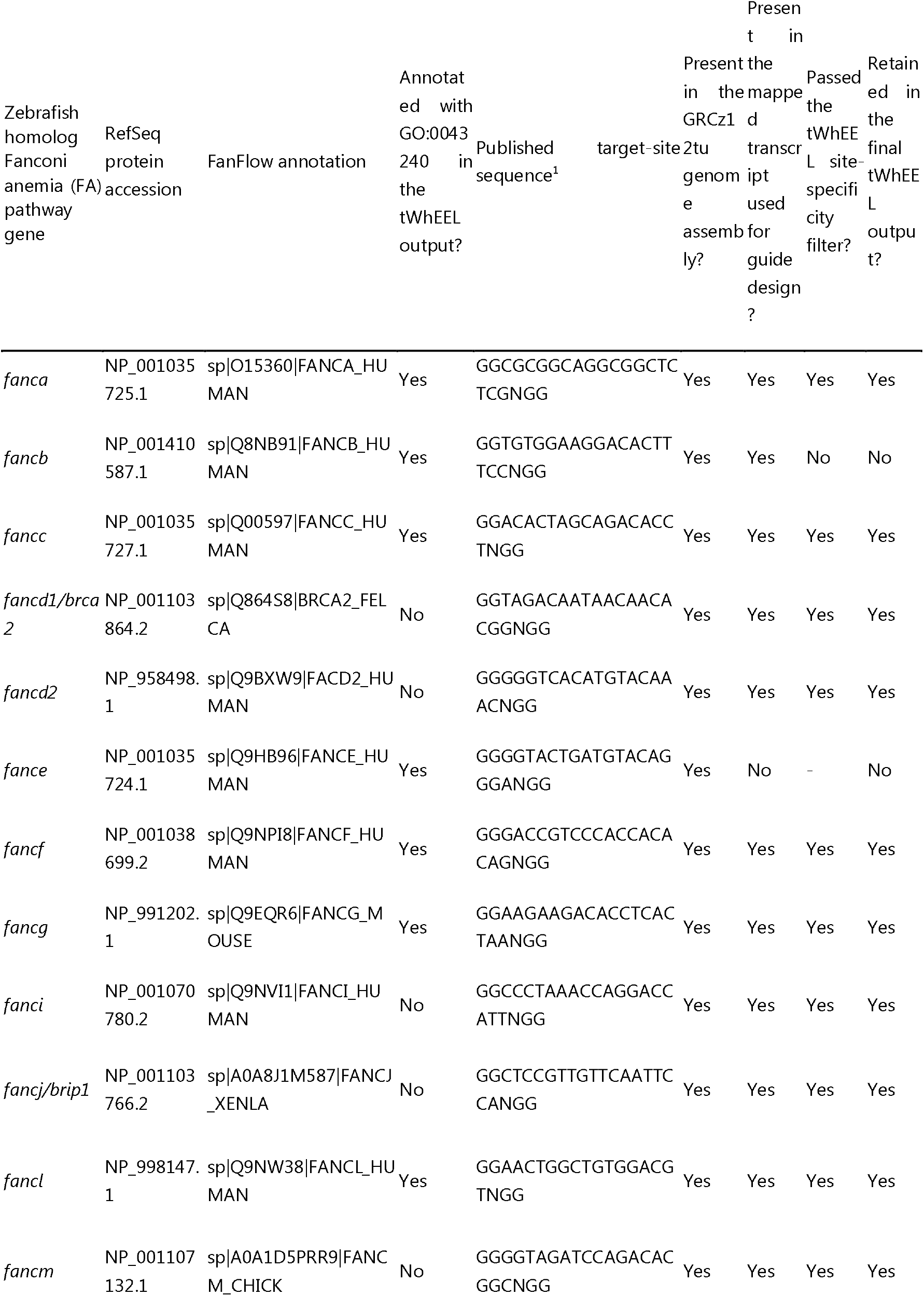
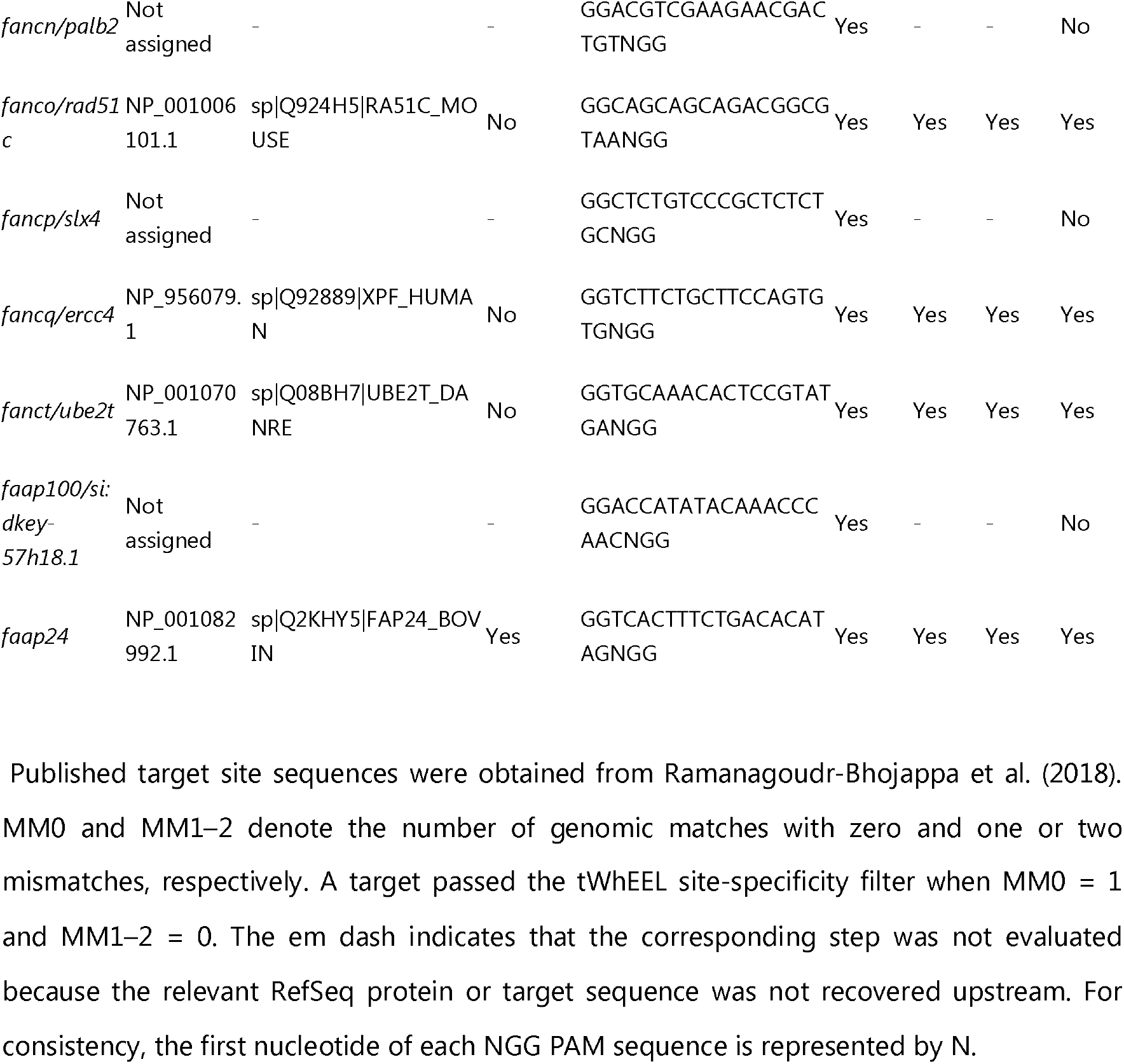
Zebrafish Fanconi anemia pathway case-study summary.

## RESULTS

### tWhEEL integrates transcriptome annotation, redundancy grouping, and guide design

The tWhEEL workflow starts with transcriptome sequences and optional genome/protein resources, and produces annotated guide RNA design reports (Figure 1A). First, protein sequences are predicted from the transcriptome or read from an existing protein FASTA file. Second, homology-based functional annotation assigns query proteins to reference proteins. Third, the queries are grouped into single-copy-like and multicopy/transcript-variant categories. Fourth, the CRISPR guide design is performed for single groups and aligned with multigroup consensus sequences. Finally, annotation-level targetability is summarized using KFI calculations (Figure 1B, Supplementary Figure 1). tWhEEL calculates the GO-KFI for GO terms and PW-KFI for pathways with available annotation resources.

Existing guide RNA design tools typically return candidate target sites in the genome sequence, whereas effective knockout design requires knowledge of whether a whole biological function, GO term, or pathway can be knocked out in an interpretable manner. tWhEEL connects the layers of both guide design and gene group functionality by carrying gene-group identifiers from annotation through the guide design into the final report. The HTML output provides a browser-based interface for sorting annotations using KFI, inspecting gene groups, and downloading guide RNA candidates.

### KFI provides an interpretable annotation-level measure of knockout feasibility

GO-KFI was calculated as the proportion of gene groups assigned to a GO term with at least one guide candidate. This definition makes the score transparent; a GO-KFI of 1 indicates that every gene group assigned to the GO term is targetable using the chosen sgRNAs, whereas a GO-KFI of 0 indicates that no annotated gene group is currently targetable. Intermediate values identify GO terms for which only a subset of annotated gene groups can be targeted.

We performed GO-KFI calculations using the Ensembl Fungi cDNA and protein sequences of Saccharomyces cerevisiae CEN.PK113-7D. In the yeast CEN.PK113-7D analysis, nearly all GO terms were fully targetable (5,622 of 5,655 terms; 99.4%), with 36.3% classified as fully and simply targetable and 63.1% as fully and multiply targetable. The four-class representation further separated fully targetable terms into simple and multiple classes (Figure 1C). "Fully and simply targetable (F & S)" GO terms, which accounted for 36.3%, have a single annotated gene group and at least one guide RNA, making them favorable candidates for straightforward knockout experiments. "Fully and multiply targetable (F & M)" GO terms, which accounted for 63.1%, have multiple annotated groups, all of which are targetable; these may still be experimentally feasible but may require multiplex editing or careful interpretation. "Partially targetable (P)" terms, which reached 0.6%, flag incomplete target coverage within the hit gene groups, and "not targetable" terms (N) mean that no sgRNA targets a gene in the functional gene groups. The only GO term classified as not targetable was lipoyl-dependent peroxiredoxin activity (GO:0160319). The associated annotation-based design unit contained one yeast query protein whose best hit was a bacterial AhpD reference protein; no guide was retained for this unit. Overall, 5,654 of 5,655 GO terms included at least one design unit with a retained guide, and 5,622 (99.4%) had at least one retained guide for every associated design unit. These results demonstrate broad guide availability for functionally annotated design units in this yeast dataset.

The HTML report of the GO analysis provided KFI statistics and a guide design list linked to GO information and functional gene groups (Figure 2A). Users can also refer to the gene name, the related functional gene group, and the guide design count (Figure 2B). This report provided a guide profile targeting a query classified into a functional gene group (Figure 2C). The tWhEEL GO analysis counts GO terms flagged as “Fully targetable” by GO parent terms, which are general ontology terms for categorizing biological function sets and sorts them in descending order (Figure 2D). GO-based phenotyping is difficult because the definitions of multiple GO terms overlap. In this case, counting GO terms by parent term, which represents a broader set of biological functions, could be helpful for estimating the feasibility of KO phenotyping. The tWhEEL function was designed to help researchers find obvious KO phenotypes based on the high “Fully targetable” count on a parent term.

**Figure 2:**
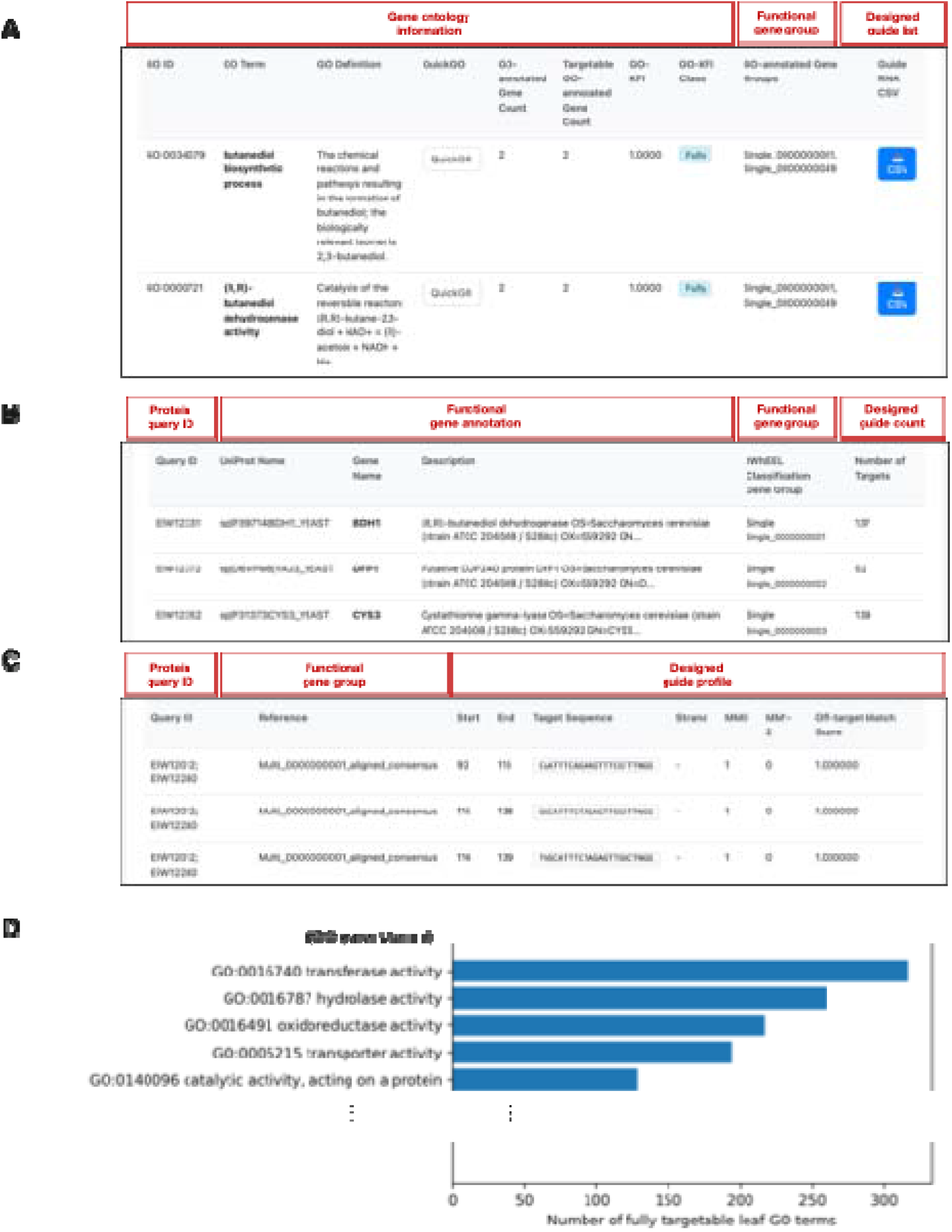
Four report views show GO-term summaries, annotated protein-query groups, guide-level records and a GO Slim summary of fully targetable terms. (A) GO-term overview linking functional annotations, contributing gene groups and downloadable guide lists. (B) Annotation records with gene-group classifications and guide counts. (C) Guide-level records with reference sequences, coordinates, strand, and mismatch counts. (D) Summary of fully targetable GO terms by GO Slim parent category.

### GO-KFI captures differences among model and nonmodel datasets

GO-KFI distributions differed among the zebrafish RefSeq query protein sets (Supplementary Table 2). The RefSeq NCBI Protein (NP)-only analysis assessed 15,698 GO terms, of which 31.2% were fully and simply targetable, 67.8% were fully and multiply targetable, 1.0% were partially targetable, and 0.0% were not targetable after rounding. Including Predicted Protein model (XP)-prefixed proteins increased the number of assessed GO terms to 17,958 and shifted their proportions to 24.6%, 71.3%, 3.9%, and 0.3%, respectively. The overall proportion of the fully targetable terms decreased from 99.0% to 95.9%. Both analyses used the same UniProtKB/Swiss-Prot annotation database and GRCz12tu genomic specificity reference, illustrating how query protein selection changes the composition of annotated groups and the resulting KFI distribution; however, it does not establish a change in the biological knockout feasibility of zebrafish. This comparison emphasizes the importance of defining the transcript and protein inputs when interpreting function-level targetability. Therefore, we used a common input processing workflow for cross-species comparisons.

To compare species under a common input-processing workflow, we performed *de novo* ORF prediction on transcriptome assemblies from the model organisms *Homo sapiens* (GenBank: GLNK00000000.1), *Danio rerio* (GenBank: GFIL00000000.1), *Drosophila melanogaster* (GenBank: GHHI00000000.1), *Arabidopsis thaliana* (GenBank: GKIF00000000.1), and *Saccharomyces cerevisiae* Y12 (GenBank: GFJR00000000.1), as well as the nonmodel organisms *Nannochloropsis gaditana* (GenBank: GAGR00000000.1), *Mimachlamys varia* (GenBank: GGGO00000000.1), and *Entoria okinawaensis* (GenBank: ICSG00000000.1) with the common default settings. The resulting predicted proteomes were subjected to functional annotation and BUSCO analysis, and the GO-KFI values were calculated (Table 1).

The total number of GO hits assigned by functional annotation was 17,858 for *Homo sapiens* and 17,664 for *Danio rerio*, placing these species among those with the highest values. In contrast, *Saccharomyces cerevisiae* Y12 strain had only 5,884 GO hits, which was the lowest number among the species examined. The nonmodel organism *Mimachlamys varia* had 17,116 GO hits, which was comparable to that of humans.

With respect to GO-KFI, *Nannochloropsis gaditana* showed the highest proportion of fully and simply targetable GOs—GO terms represented by a single annotation-based design unit with at least one retained guide—at 46.6%. The lowest proportion was observed in *Arabidopsis thaliana* (18.8 %). In *A. thaliana*, partially targetable GO terms, for which guides were available for only a subset of annotated gene groups, accounted for 54.2%. The KFI can alert that other genome-editing platforms, rather than simple SpCas9 mutagenesis, may be worth considering for genome-editing studies in such cases. *Arabidopsis thaliana* also had the highest proportion of not-targetable GOs at 15.8%; these were GOs for which associated genes had been identified but could not be targeted based on the TSA dataset analyzed in this study. In the *Arabidopsis thaliana* GKIF01 dataset, 4,234 of the 14,146 GO terms were fully targetable (29.9%), including 2,663 fully and simply targetable terms (18.8%), whereas 7,671 and 2,241 terms were partially targetable (54.2%) and not targetable (15.8%), respectively. Candidate sites were detected in all 37,567 annotated design units; however, only 24,705 units (65.8%) retained at least one guide after site-specific filtering. Retention was observed in 11,195 of 15,864 single units (70.6%) and 13,510 of 21,703 multiunits (62.2%). Among the 12,862 units without retained guides, 12,243 (95.2%) had no candidate with exactly one perfect match to the specificity reference (see Discussion).

tWhEEL also summarizes targetable biological functions by mapping fully targetable GO terms to GOSlim parent categories. We listed the top ten parent GO terms (Supplementary Table 3), and found five common terms (catalytic activity, acting on a protein (GO:0140096); hydrolase activity (GO:0016787); oxidoreductase activity (GO:0016491); transferase activity (GO:0016740); transporter activity (GO:0005215)) among the eight TSA datasets. One GO term, carbohydrate derivative metabolic process (GO:1901135), was uniquely ranked within the top ten in the Drosophila melanogaster (GenBank: GHHI00000000.1) dataset. This broad GO category encompassed biologically important pathways in insects, including nucleotide-sugar metabolism, glycosylation, and chitin biosynthesis, which may be closely associated with development and exoskeletal formation. Three GO terms, catalytic activity acting on RNA (GO:0140098), protein-containing complex assembly (GO:0065003), and nucleobase-containing small molecule metabolic processes (GO:0055086), were uniquely ranked within the top ten in the Saccharomyces cerevisiae Y12 (GenBank: GFJR00000000.1) dataset.

GO-KFI and the associated GO information can thus facilitate the estimation of an organism’s amenability to genome editing associated with a biological function described in GO terms based on its transcriptomic information. KFI therefore helps researchers assess whether an organism is suitable for investigating a biological question before investing in de novo genome sequencing.

### A zebrafish Fanconi anemia pathway case study recovers published CRISPR guide designs

To evaluate whether the tWhEEL workflow can recover plausible target designs in a biologically interpretable gene cluster, we ran tWhEEL with the GRCz12tu genome and Known RefSeq protein annotation (with NP_ accession prefix) and then compared the tWhEEL output with a published multiplex CRISPR-Cas9 knockout study of 19 FA pathway genes in zebrafish. tWhEEL recovered the target site designs, matching a published study for 14 of the 19 target genes (Table 2). Eight of the 19 target genes were annotated using the GO term “Fanconi anemia nuclear complex” (GO:0043240).

The five unrecovered published designs were explained by query protein representation, transcript–genome sequence differences, or sequence-specificity filters. The three genes *fancn/palb2*, *fancp/slx4*, and *faap100* were not detected in the Known RefSeq protein annotation set because their relevant records were categorized as computationally predicted proteins (with the XP_ accession prefix). Query protein set selection can change both the KFI values and apparent gene recovery. The guide sequence for fance was also not detected with tWhEEL because the sequence of fance-annotated transcripts differed from the GRCz12tu genomic sequence at protospacer position 4. The public GFF annotation file from NCBI notes that the RefSeq transcript of fance contains 12 substitutions compared to this genomic sequence. The fance example illustrates how transcript-derived guide discovery exposes sequence differences between a transcript record and the corresponding reference genome, allowing these differences to be considered during experimental design. Lastly, the guide sequence targeting the fancb gene was identified during guide discovery but was excluded by the sequence-specificity filter. Thus, tWhEEL retained guide candidates with one perfect match and no one- or two-mismatched matches in the selected reference.

### PW-KFI enables pathway-level guide inspection when pathway-gene identifiers are available

The pathway module extends the same KFI concept to pathway gene sets using WikiPathways GPML files (Figure 3A). In the yeast demonstration, the tWhEEL-parsed pathway diagrams (Figure 3B) matched pathway genes to SGD identifiers, calculated the PW-KFI, and generated an interactive report. Among the 94 pathways reported for CEN.PK113-7D, 71 (75.5%) were fully targetable, and 23 (24.5%) were partially targetable. Histidine biosynthesis (WP514) retained guides for all seven evaluable genes and sulfur amino acid biosynthesis (WP7) retained guides for all 14 genes, yielding PW-KFI = 1 for both pathways. The 10 gene nodes in the WP514 diagram correspond to seven distinct genes, illustrating how repeated pathway nodes were consolidated before the KFI calculation. In contrast, the meiotic recombination gene set (WP377), shown in Figure 3A, retained guides for 15 of the 16 evaluable genes (PW-KFI = 0.9375). These examples connect the pathway-level feasibility to genes and guide candidates to support an experimental plan. In this study, pathways were sorted by total gene count, guide-covered gene count, PW-KFI, and coverage rate. Pathway diagrams displayed guide availability at the gene level, and gene-detail models provided guide sequences, coordinates, strands, and off-target summary values (Figure 3C). This pathway-level view is useful because biological perturbations are often designed around pathway hypotheses, rather than isolated GO terms.

**Figure 3:**
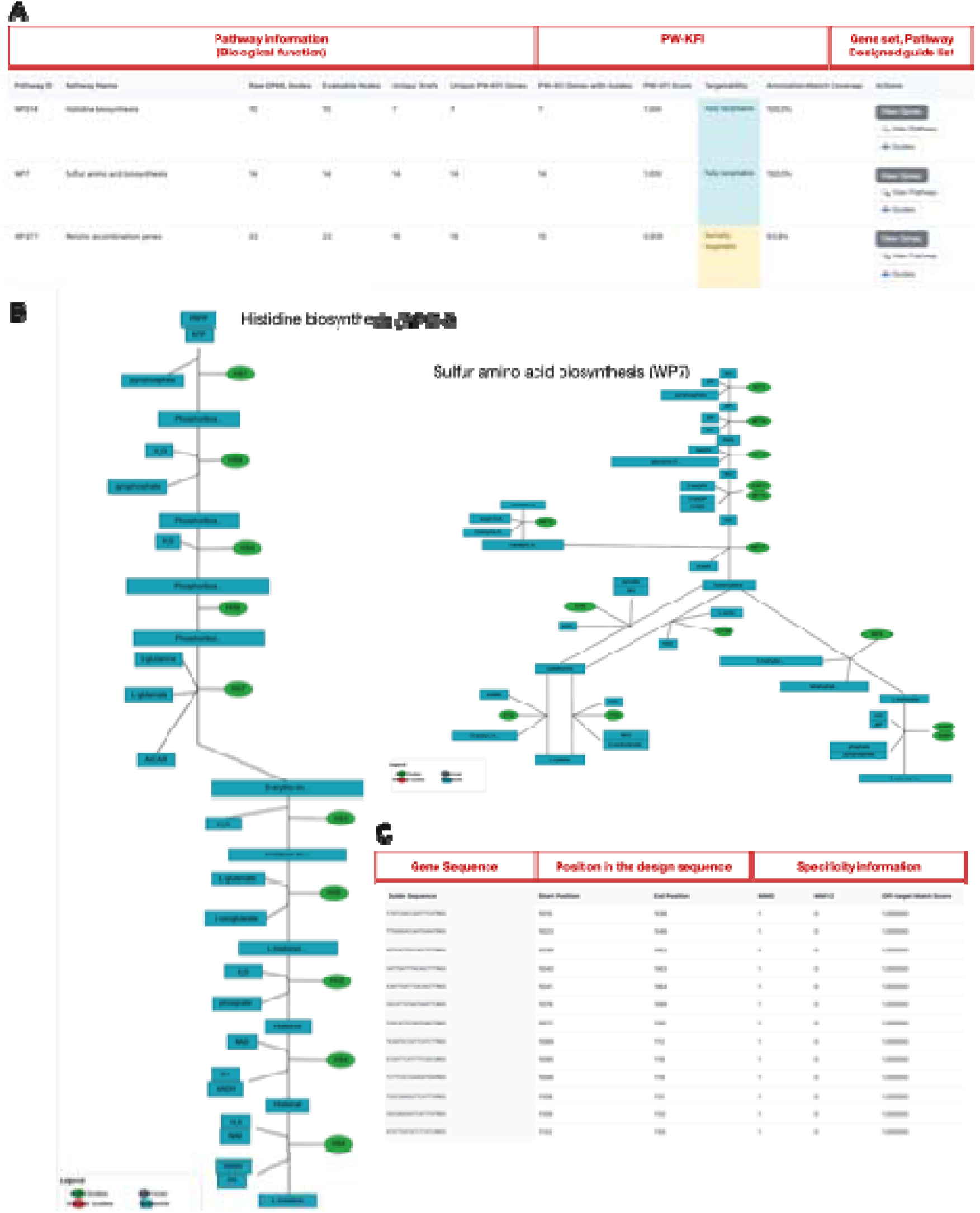
Interactive pathway-KFI report. (A) Pathways are listed with total genes, genes with guides, PW-KFI scores, targetability classes and coverage. (B) GPML-derived pathway diagrams highlight genes by guide availability. (C) Gene-detail views provide guide sequences, coordinates and off-target summary information. Alt text: Screenshots of a pathway table, a colored pathway diagram and a guide RNA detail modal.

Together, these results demonstrate how the tWhEEL pipeline assesses the feasibility of gene knockout in user-selected species, strains, and tissues and provides a user-friendly report that includes sgRNA target data, gene annotations, GO information, and pathway information. The pipeline facilitates the design of gene knockout studies across diverse organisms.

## DISCUSSION

tWhEEL reframes CRISPR guide design as an annotation-aware feasibility problem. Rather than asking only whether a guide RNA can be designed for a sequence, tWhEEL asks whether annotated gene groups underlying a function or pathway can be targeted under a specific set of assumptions. Targetability is important in reverse genetics, where gene duplication, transcript diversity, and reference database choices can determine whether a knockout experiment is interpretable.

The workflow is particularly relevant for nonmodel organisms. Many emerging organisms have transcriptome assemblies and functional annotations before the development of chromosome-level genomes, curated gene models, or organism-specific pathway databases. tWhEEL can operate in this intermediate state using a transcriptome for protein prediction, and when necessary, for guide design. The resulting KFI values should not be interpreted as exhaustive genome-wide off-target safety assessments in the absence of a genome assembly. These summaries can help identify organisms and biological functions suitable for reverse genetic investigations before investing in *de novo* genome sequencing.

The assembled transcriptome rather than whole genome was used as a specificity reference to apply a common guide selection procedure across TSA datasets. Under these conditions, GO-KFI reflects the availability of guides that uniquely match the represented transcript sequences in the presence of alternative transcripts, multicopy genes, and closely related paralogs. We used *the Arabidopsis thaliana* TSA dataset, GKIF00000000.1, as one of the benchmark input datasets. The GKIF00000000.1 TSA dataset was built as a comprehensive transcript reference dataset (AtRTD3) for *Arabidopsis thaliana* using various tissues and developmental stages. Previous studies have reported alternative splicing in approximately 61% of multiexonic genes in A. thaliana under normal growth conditions (38). Moreover, a meta-analysis of 206 publicly available RNA-seq samples using the AtRTD2-QUASI reference transcriptome identified differential transcript usage in 8,148 genes (39). The finding that low GO-KFI in the *Arabidopsis thaliana* TSA dataset, GKIF00000000.1, arose during sequence-specificity filtering, rather than from the absence of candidate sites, may reflect transcriptome complexity. Guide selection for *A. thaliana* can, therefore, consider the transcripts expressed in the tissue and developmental stage relevant to the phenotype of interest, together with specificity against the genomic reference.

The zebrafish RefSeq comparison showed how the query protein selection shaped the functional groups represented in the KFI analysis. Including XP-prefixed records adds predicted protein models and changes the distribution of targetability classes, whereas NP-only inputs omitted genes represented exclusively by XP-prefixed records. Recording the query-set composition alongside the annotation database provenance therefore makes these comparisons interpretable.

This study has several limitations. The application of KFI depends on the available sequences and annotation resources. Input selection and functional annotation determine the groups represented in GO-KFI, whereas pathway analysis requires identifiers that can be resolved into evaluable genes. Sequence-specificity filtering uses supplied genome or transcriptome references. For the experimental implementation, users select guide candidates that are appropriate for the intended loss-of-function strategy and assess their activities. By linking each functional summary to its underlying sequences, groups, and guide records, tWhEEL makes these choices accessible during experimental planning.

We note several points with regard to the interpretation of the results: (i) KFI quantifies the feasibility of the genome editing design under the selected sequence inputs, annotations, and filtering criteria. This does not guarantee that each guide RNA has high editing efficiency. he resulting guide candidates also require experimental evaluation of guide activity. (ii) Specificity estimates depend on the reference sequence collection. Transcriptome-only searches omit unrepresented genomic sequences and count shared regions of different transcript records as multiple matches. Genome-backed searches, provided by tWhEEL’s optional function, use a broader reference; however, the requirement of tWhEEL for one perfect match and no one- or two-mismatch matches does not assess all potential off-target mechanisms or verify the intended locus and exon. Therefore, experimental designs require independent off-target prediction using additional software (40, 41) or sequencing analysis using whole-genome samples (42, 43). (iii) The coverage of functional gene groups is also conditional on annotation and identifier compatibility. The GO-KFI excludes unannotated contributors to functions. PW-KFI includes evaluable genes lacking an annotation match, but excludes nodes whose identifiers cannot be evaluated under the configured rules. Consequently, a PW-KFI of 1 indicates evaluable pathway genes that may coexist with unassessed nodes in the original pathway. Annotation coverage and identifier audits should also be considered along with the score. Cross-dataset comparisons require a consistent input selection, specific references, and annotation resources.

For experimental use, sgRNAs reported by tWhEEL should be treated as candidates rather than final designs. Delivery vector architecture, promoter choice, Cas nuclease, PAM preference, target exon position, isoform structure, multiplexing strategy, and genotyping plan should be considered. For loss-of-function experiments, upstream coding exons are often preferred; however, the best target site depends on the locus and the intended assay. tWhEEL was designed to expose these choices by linking each functional category to the underlying gene groups and guide RNA records.

tWhEEL provides a transcript-aware, annotation-centered framework to evaluate the feasibility of CRISPR knockouts. By integrating transcriptome-derived gene prediction, homology-based functional annotation, redundancy-aware grouping, guide RNA design, and GO/pathway-level KFI scoring, this workflow helps researchers move from sequence-level guide candidates to function-level experimental planning. We demonstrated that KFI provided a quantitative measure of genome-editing targetability among eight organisms and that the tWhEEL guide design identified 14 of 19 published target sites from a previous experimental study. We believe that the tWhEEL and KFI calculation frameworks are useful for comparing reverse genetic tractability and for prioritizing focusing biological functions for model and nonmodel organisms in reverse genetics research.

## ACKNOWLEDGEMENTS

We thank the members of the Laboratory of Bio-DX at the Genome Editing Innovation Center and the Laboratory of Genome Informatics at the Graduate School of Integrated Sciences for Life, Hiroshima University for helpful discussions.

OpenAI Codex was used to assist with the software and analysis-script development, preparation of figure and table drafts, English-language editing, and checks for consistency between the manuscript, software implementation, and analysis outputs. The authors reviewed and revised the AI-assisted materials and assume responsibility for the final content.

## AUTHOR CONTRIBUTIONS

K.N.: Conceptualization, Software, Methodology, Formal analysis, Visualization, Writing – original draft, Data curation, Validation, Investigation. H.B.: Conceptualization, Supervision, Funding acquisition, Writing – review & editing.

## SUPPLEMENTARY DATA

Supplementary Figure 1 illustrates transcript grouping, consensus generation, sequence-specificity filtering, and the counting units used to calculate GO-KFI and PW-KFI. Supplementary Table 1 lists reference resources and BUSCO lineage datasets, including version information and checksums. Supplementary Table 2 compares GO-KFI targetability categories between the zebrafish NP-only and NP+XP query protein sets. Supplementary Table 3 presents the top ten GO Slim parent categories of fully targetable GO terms for each of the eight transcriptome assemblies.

## DATA AVAILABILITY

The tWhEEL source code, Docker files, configuration templates, and report-generation assets are available at https://github.com/KazukiNakamae/tWhEEL under the MIT License. The released software version is v1.3.5. The version was archived at Zenodo (https://doi.org/10.5281/zenodo.23660612). Input-resource accession lists, checksums, and numerical results are presented in Tables 1 and 2 and Supplementary Tables 2 and 3. The used dataset and outputs are available from Figshare at https://figshare.com/account/articles/34786824. The reference resource versions and BUSCO lineage dataset are listed in Supplementary Table 1.

