## Supplementary Information for "tWhEEL: a transcript-aware workflow for quantifying CRISPR knockout feasibility using transcriptome assembly"

**Contents**

1. Supplementary Figures . . . . . . . . . . . . . . . . . . . . . . . . . . . . . . . . . . . . . . . . . . . . . . . . . .. . . . 2
2. Supplementary Table Legends . . . . . . . . . . . . . . . . . . . . . . . . . . . . . . . . . . . . . . . . . . . .. . . . 4

**1. Supplementary Figures**

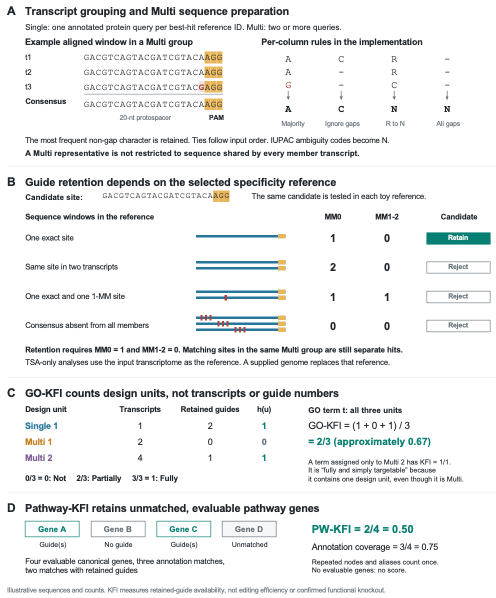

Supplementary Figure 1. From transcript sequences to KFI counting units.

Illustrative examples explain the tWhEEL implementation; panels are independent examples that do not represent biological benchmark observations.

(A) Annotated protein queries sharing a best-hit reference identifier form a single group when there is one query, and a multi-group when there are two or more. For clarity, the examples show an annotated protein query per transcript. The aligned multiwindow illustrates the selection of the most frequent nongap characters in each column. Ties follow an aligned record order and all-gap columns become N. Supported IUPAC ambiguity codes other than N are converted to N before guide discovery. The representative sequence does not need to be shared by every member transcript; the displayed protospacer–PAM window is a candidate site, and there is no evidence that it will be retained or target all members.

(B) The same synthetic 20-nt protospacer as AGG PAM was assessed against four separate toy-specificity references. CrisFlash reports one exact match alone (MM0 = 1, MM1–2 = 0), duplicate exact matches in two transcript records (2, 0), one exact match plus one single-mismatch match (1, 1), and no match within two mismatches (0, 0). In the last example, the three-member protospacers differ from their representative consensus in nonoverlapping sets of three positions. Only the first candidate satisfied the retention criteria. The hit counts were verified by running CrisFlash on synthetic references and applying a tWhEEL filter. In the transcriptome-reference mode, repeated sites in separate transcript records remain as separate matches, even within one Multi group. When a genome is supplied, the matches are instead counted in that genomic reference. Sequence windows are shown from 5` to 3`, with flanking bases omitted; red marks indicate substitutions relative to the candidate protospacer.

(C) GO-KFI uses distinct annotated design units, not transcript counts or the sum of retained guides. The three units contain 1, 2, and 4 transcripts and retain 2, 0, and 1 guides, respectively, yielding 2/3. The stipulated guide counts are independent of the sequence examples in panels A and B, and the guides may differ between units. A GO term assigned only to the guide-positive Multi 2 unit has a KFI = 1/1 and is classified as fully and simply targetable because it contains one design unit. This subclass does not imply that the unit is a single group or that all member transcripts are targeted.

(D) The PW-KFI uses evaluable canonical genes from external pathway definition. Of the four genes, three matched the annotation and two retained the guides, yielding PW-KFI = 2/4 = 0.50 and annotation coverage = 3/4 = 0.75. The unmatched but evaluable gene remains in the denominator and repeated nodes and aliases that resolve the same gene contribute once. Nonevaluable identifiers were excluded and pathways without evaluable genes received no score. Both KFI measures describe retained-guide availability, not editing efficiency or experimentally established knockout feasibility.

**2. Supplementary Table Legends**

Supplementary Table 1. Reference resources and BUSCO lineage datasets used in the tWhEEL analyses. File names, release or version identifiers, release dates, retrieval information, SHA-256 checksums, provenance, and source URLs are provided for UniProtKB/Swiss-Prot, SGD reference proteins, GO ontology, generic GO Slim, and eight BUSCO lineage datasets. Checksums refer to the files specified in the “File used” column; BUSCO lineage checksums cover `dataset.cfg`. NA: information unavailable in retained records. GO, Gene Ontology; SGD, Saccharomyces Genome Database; BUSCO, Benchmarking Universal Single-Copy Orthologs; ORF, open reading frame; TSA, transcriptome shotgun assembly; JST, Japan Standard Time. NP-only and NP+XP denote the RefSeq query sets containing NP-prefixed records alone or both NP- and XP-prefixed records, respectively.

Supplementary Table 2. GO term categories are reported as counts and percentages of all GO terms assessed in each analysis. Both analyses used tWhEEL v1.3.4 for guide design and GO-KFI calculation, with the same UniProtKB/Swiss-Prot annotation database and the GRCz12tu genome assembly as the specificity reference. Their HTML reports were subsequently regenerated using tWhEEL v1.3.5 without repeating guide design or GO-KFI calculation. Fully and simply targetable terms had one annotated design unit with at least one retained guide. Fully and multiply targetable terms have multiple annotated design units, each with at least one retained guide. Partially targetable terms have guides for some but not all units, whereas not-targetable terms have no retained guides. The retained guides satisfied MM0 = 1 and MM1–2 = 0. The percentages are rounded to one decimal place and may not sum up to 100%. The NP-only non-targetable category contained three GO terms (0.0191%), displayed as 0.0% after rounding.

Supplementary Table 3. Top 10 GO Slim parent categories of fully targetable GO terms across eight transcriptome assemblies. GO terms with GO-KFI = 1 were mapped to generic GO-Slim parent categories for each dataset. Each mapped term was assigned to one category. Categories were ranked by the number of assigned fully targetable GO terms, and the ten highest-ranked categories are reported with their GO identifiers, descriptions, and counts. The counts represent GO terms rather than genes, transcripts, or guides. GO, Gene Ontology; KFI, knockout feasibility index; TSA, Transcriptome Shotgun Assembly.
